# Allotypic variation in antigen processing controls antigenic peptide generation from SARS-CoV-2 S1 Spike Glycoprotein

**DOI:** 10.1101/2021.07.03.450989

**Authors:** George Stamatakis, Martina Samiotaki, Ioannis Temponeras, George Panayotou, Efstratios Stratikos

## Abstract

Population genetic variability in immune system genes can often underlie variability in immune responses to pathogens. Cytotoxic T-lymphocytes are emerging as critical determinants of both SARS-CoV-2 infection severity and long-term immunity, either after recovery or vaccination. A hallmark of COVID-19 is its highly variable severity and breadth of immune responses between individuals. To address the underlying mechanisms behind this phenomenon we analyzed the proteolytic processing of S1 spike glycoprotein precursor antigenic peptides by 10 common allotypes of ER aminopeptidase 1 (ERAP1), a polymorphic intracellular enzyme that can regulate cytotoxic T-lymphocyte responses by generating or destroying antigenic peptides. We utilized a systematic proteomic approach that allows the concurrent analysis of hundreds of trimming reactions in parallel, thus better emulating antigen processing in the cell. While all ERAP1 allotypes were capable of producing optimal ligands for MHC class I molecules, including known SARS-CoV-2 epitopes, they presented significant differences in peptide sequences produced, suggesting allotype-dependent sequence biases. Allotype 10, previously suggested to be enzymatically deficient, was rather found to be functionally distinct from other allotypes. Our findings suggest that common ERAP1 allotypes can be a major source of heterogeneity in antigen processing and through this mechanism contribute to variable immune responses to COVID-19.

## INTRODUCTION

Immune responses to Severe Acute Respiratory Syndrome Coronavirus 2 (SARS-CoV-2), the pathogen behind coronavirus disease 19 (COVID-19), play critical roles in disease pathophysiology^1,2^. Appropriate innate and adaptive immune responses are necessary for viral clearance, while aberrant uncontrolled responses have a major impact on mortality^3,4^. Furthermore, long-term immunity after infection or vaccination is of critical importance for ending the current pandemic ^5^. Although initial analyses focused on antibody dependent immunity, T-cell mediated immunity appears to be important for both viral clearance and for long-term immunity, especially against emerging virus variants ^6–8^. As a result, detailed knowledge of how SARS-CoV-2 epitopes are generated and selected is crucial for both allowing a better understanding of long-term anti-viral immune responses by individuals and for the optimization of vaccines against virus variants^9^.

COVID-19 is characterized by significant variability in disease severity amongst individuals ^10^. As a result, several studies have focused in determining genetic predispositions to COVID-19 in an effort to establish better preventive and treatment measures ^11,12^. Indeed, rare mutations in components of the innate immune response and in particular the interferon I pathway can predispose individuals to severe COVID-19 ^13,14^. In addition, the role of common polymorphic variation in components of adaptive immunity, such as Human Leukocyte Antigen(HLA)-alleles, has been emerging an a major factor ^15–17^.

HLA molecules (also called Major Histocompatibility Complex molecules, MHC) bind small peptides generated intracellularly from antigenic proteins of pathogens and present them on the cell surface. The HLA class I - peptide complexes interact with specialized receptors on CD8^+^ T cells and successful recognition indicates an infected cell and initiates a molecular response that leads to cell lysis ^18^. Several enzymes play roles in generating peptide ligands for HLA molecules. ER aminopeptidase 1 (ERAP1) is an ER-resident aminopeptidase that trims precursors of antigenic peptides to optimize them for binding onto HLA ^19^. Appropriate of aberrant trimming of antigenic peptide precursors by ERAP1 can alter the repertoire of peptides available for presentation by HLA and indirectly regulate adaptive immune responses ^20^. Thus, the generation of antigenic epitopes by ERAP1 can be critical for immune evasion by viruses ^21,22^.

HLA molecules are highly polymorphic with thousands of different alleles discovered to date ^23^. This polymorphic variation allows for the binding of a vast variety of peptide sequences, ensuring sufficient presentation of epitopes from unknown pathogens on a population but not necessarily an individual level. Thus, HLA variability can underlie differences in immune responses between individuals. ERAP1 is also polymorphic, and coding single nucleotide polymorphisms (SNPs) in the ERAP1 gene associate with predisposition to autoimmunity or cancer often in epistasis with HLA alleles ^24^ and can help shape the cellular immunopeptidome ^25^. ERAP1 SNPs are found in the population in limited combinations that define specific allotypes and have functional consequences in peptide trimming ^26,27^. The exact interplay of ERAP1 and HLA polymorphic variation in determining antigen presentation is currently a subject of active research^21,27–29^.

Generation and loading of antigenic peptides on HLA occur inside the ER, where thousands of different peptides compete with each other. While *in vitro* enzymatic analysis of ERAP1 function has provided significant insights on the mechanism of the enzyme ^30,31^, complex interactions between substrates and ERAP1’s large substrate-binding cavity result in a complex landscape of poorly understood specificity determinants ^30,32^. To address these complexities we previously devised a proteomic approach to concurrently analyze hundreds of trimming reactions and compared the function of different antigen processing enzymes ^33^. Here, we applied this approach to analyze the relative effect of the 10 most common ERAP1 allotypes (covering 99.9% of the European population) in processing antigenic peptide precursors derived from the S1 spike glycoprotein of SARS-CoV-2. Our analysis suggests that ERAP1 allotypes carry significant sequence biases and generate sufficiently different peptide repertoires, which, in combination with HLA peptide binding specificity, can underlie differences in adaptive immune responses that contribute to COVID-19 disease severity variability in natural populations.

## RESULTS

To explore how different ERAP1 allotypes can process S1 spike glycoprotein and generate putative antigenic peptides we utilized a library of 315 synthetic 15mers, spanning the sequence of the S1 spike glycoprotein, with 11-residue overlap between adjacent peptides. These 15mers peptides model possible antigenic peptide precursors that are generated in the cytosol and enter the ER, where they are further digested by ERAP1 to generate smaller products that bind and are presented by MHC class I. Thus, this mixture constitutes a useful tool for the systematic sampling of the entire sequence of the protein and has been used before to compare the activities of ERAP1 to other enzymes in the antigen processing pathway^33^. To digest the mixture, we used highly-pure recombinant ERAP1 protein variants corresponding to naturally-occurring allotypes. These allotypes are defined as combinations of 9 SNPs in the ERAP1 gene as shown in Table 1 and constitute the most common allotypes in humans, covering 99.9% of European population and 94.1% of the global population^27^.

**Table 1:** Amino acid composition at polymorphic positions for the 10 most common ERAP1 allotypes.

| Amino acid composition at each SNP |  |  |  |  |  |  |  |  |  |
| --- | --- | --- | --- | --- | --- | --- | --- | --- | --- |
| Allotype | 56 | 127 | 276 | 346 | 349 | 528 | 575 | 725 | 730 |
| 1 | E | P | I | G | M | K | D | R | Q |
| 2 | E | R | I | G | M | K | D | R | Q |
| 3 | E | R | I | G | M | K | D | R | E |
| 4 | E | R | I | G | M | R | D | R | E |
| 5 | E | R | I | D | M | R | D | R | E |
| 6 | E | P | I | G | M | R | D | R | E |
| 7 | K | P | I | G | M | R | D | R | E |
| 8 | E | P | M | G | M | R | D | R | E |
| 9 | E | P | M | G | M | R | N | R | E |
| 10 | E | P | I | G | V | R | N | Q | E |

Since the trimming of antigenic peptide precursors by ERAP1 is a dynamic phenomenon, the peptide pool was mixed with recombinant ERAP1 at two concentrations (100 nM and 300 nM, henceforth called low-enzyme and high-enzyme condition) so as to provide us with insight on the kinetics of the reaction. After incubation, the digestion products were analyzed by LC-MS/MS using a custom search database generated by *in silico* digestions of the full S1 spike glycoprotein sequence (UniProt ID: P0DTC2). Three biological replicates for each reaction and three replicates of a negative control reaction were performed, totaling to 66 samples, and the identified peptides were filtered for robustness of detection as described in the experimental section.

To evaluate the relative progress of each digestion we first compared the total peptide abundance present after each reaction (Figure 1). We grouped the peptides into two categories: i) 15mers, that correspond to the undigested peptides (substrates) and ii) 7-14mers that correspond to the products of the digestions. For both reaction conditions (low enzyme and high enzyme) 70-80% of peptide signal came from digestion products for most allotypes, with relatively small differences between most allotypes. About 20% of 15mers were still present after digestion for both reaction conditions, likely representing peptides that are resistant to ERAP1 trimming. Allotype 10 was slightly less efficient, with 54-62% of peptide signal originating from product peptides. Still, this difference is much less pronounced than observed in previous *in vitro* experiments using single peptides that suggested that allotype 10 can be as much as 60-fold less active for some peptides^27^. This finding suggests that although allotype 10 may be less active for some substrates, it is still able to operate with good efficiency in complex substrate mixtures.

**Figure 1:**
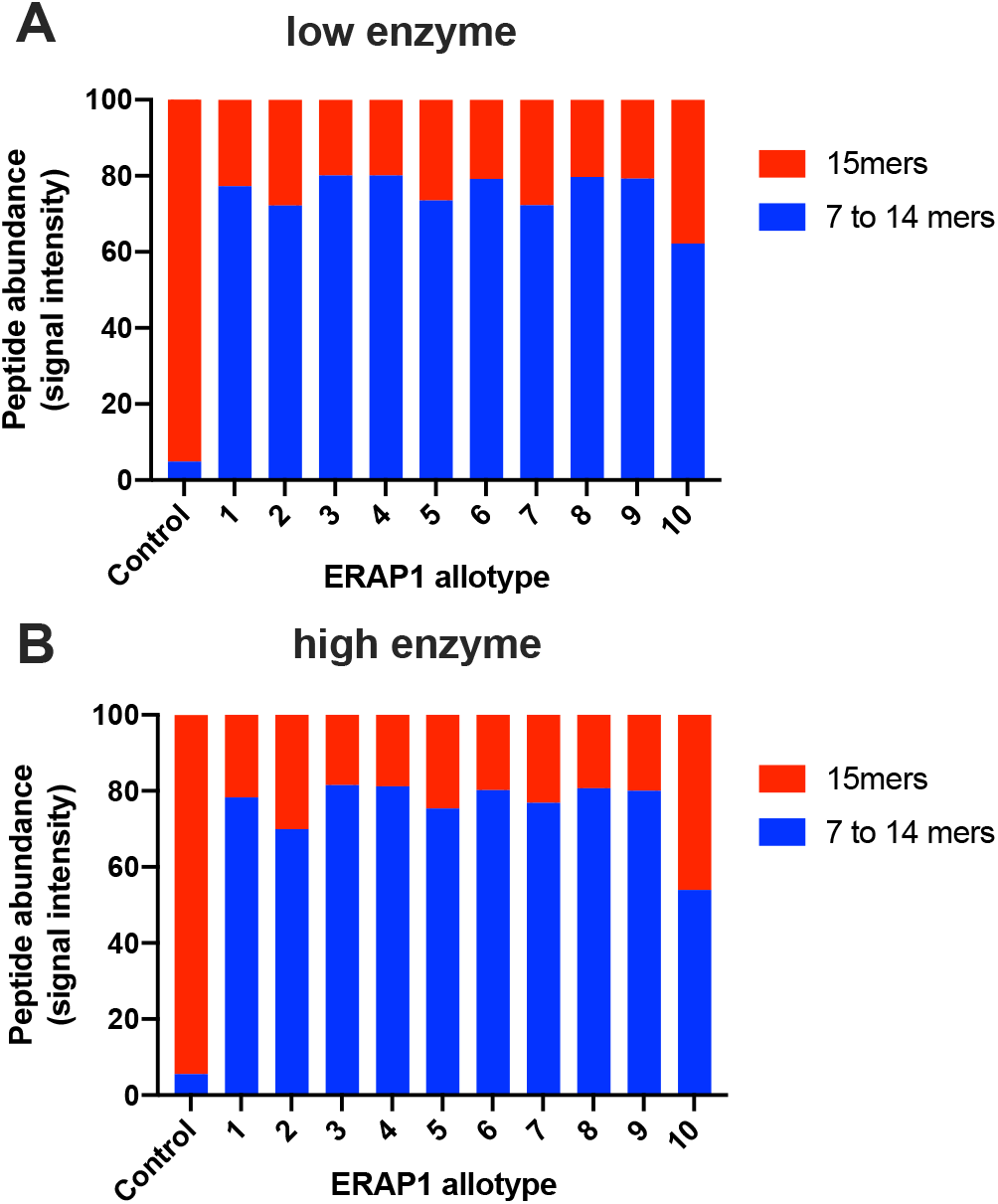
Relative peptide abundance in each sample, calculated from the total signal intensity, shown for the initial 15mer peptides and the produced 7-14mers, for both enzyme concentrations used. Control reaction was performed by incubating the peptides in the absence of enzyme.

To gain insight on the specific differences between allotypes we generated heat-map plots of all reactions and performed cluster analysis as described in the methods section (Figure 2). For both enzyme concentrations the pool of 15mer peptides was digested significantly as demonstrated by the reduction of intensity of detected 15mers (Figure 2A and 2C). Some differences were, however, evident between ERAP1 allotypes. This was most evident for allotype 10, which spared clusters of peptides, possibly due to its lower enzymatic activity as demonstrated before ^27^. Still, allotype 10 was able to efficiently trim large peptide clusters suggesting that its lower activity may be sequence specific.

**Figure 2:**
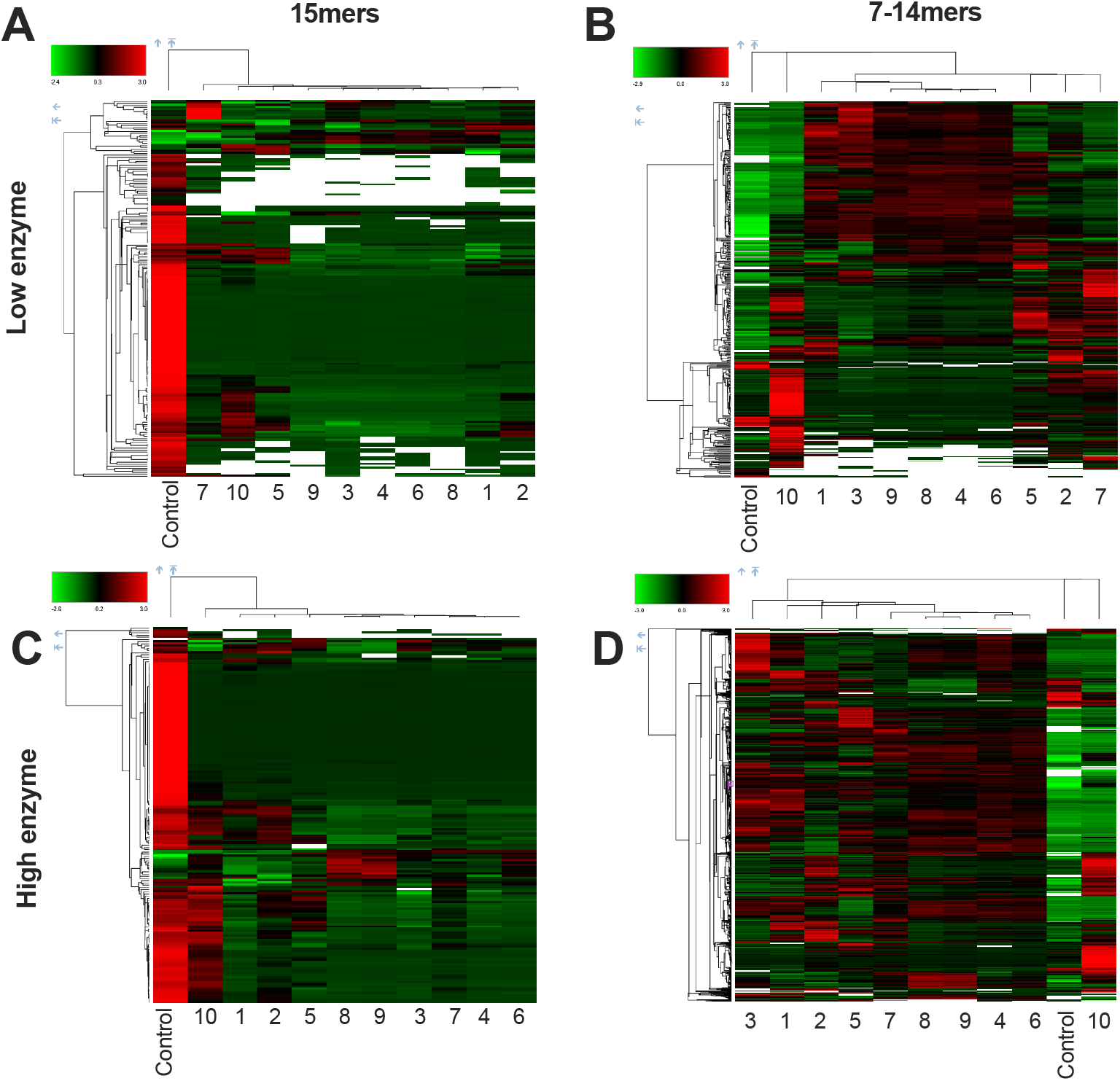
Heat-plots showing trimming of the pool of 15mer peptides (panels A and C) by all ERAP1 allotypes (allotypes 1-10) and generation of smaller peptide products 7-14 amino acids long (panels B and D) under two different conditions containing 100 nM and 300 nM enzyme respectively (low enzyme condition, panels A and B; high enzyme condition, panels C and D). Samples are organized after cluster analysis performed by Proteome Discoverer 2.5 (indicated by connecting lines). Control: peptides incubated in the absence of enzyme.

In terms of production of 7-14mers, there were again apparent differences in the pattens between allotypes (Figure 2B and 2D). At both enzyme concentrations, allotype 10 appeared to under-produce several peptides, but to also over-produce some peptide clusters. Similar, although less pronounced, phenomena were evident for other allotypes and in general, allotypes 4,6,8 and 9, appear to be the most efficient in producing a variety of different peptide products and tend to group together in the cluster analysis. Overall, while all allotypes were able to digest the 15mer substrates, significant differences in patterns are evident suggesting some differences in sequence preferences between allotypes.

Peptide length is one of the most important parameters for binding onto MHC class I molecules and ERAP1 has been shown to show significant preferences for substrate length ^30,34^. Most of the known MHCI ligands are in the range of 8-12 amino acids and the majority are 9mers. We thus analyzed the length distribution of produced peptides from each ERAP1 allele and for each of the two reaction conditions (Figure 3). For this, and following, comparisons, we restricted analysis only for peptides that were detected in replicate measurements to be statistically significant (P value < 0.05) compared to the control reaction.

**Figure 3:**
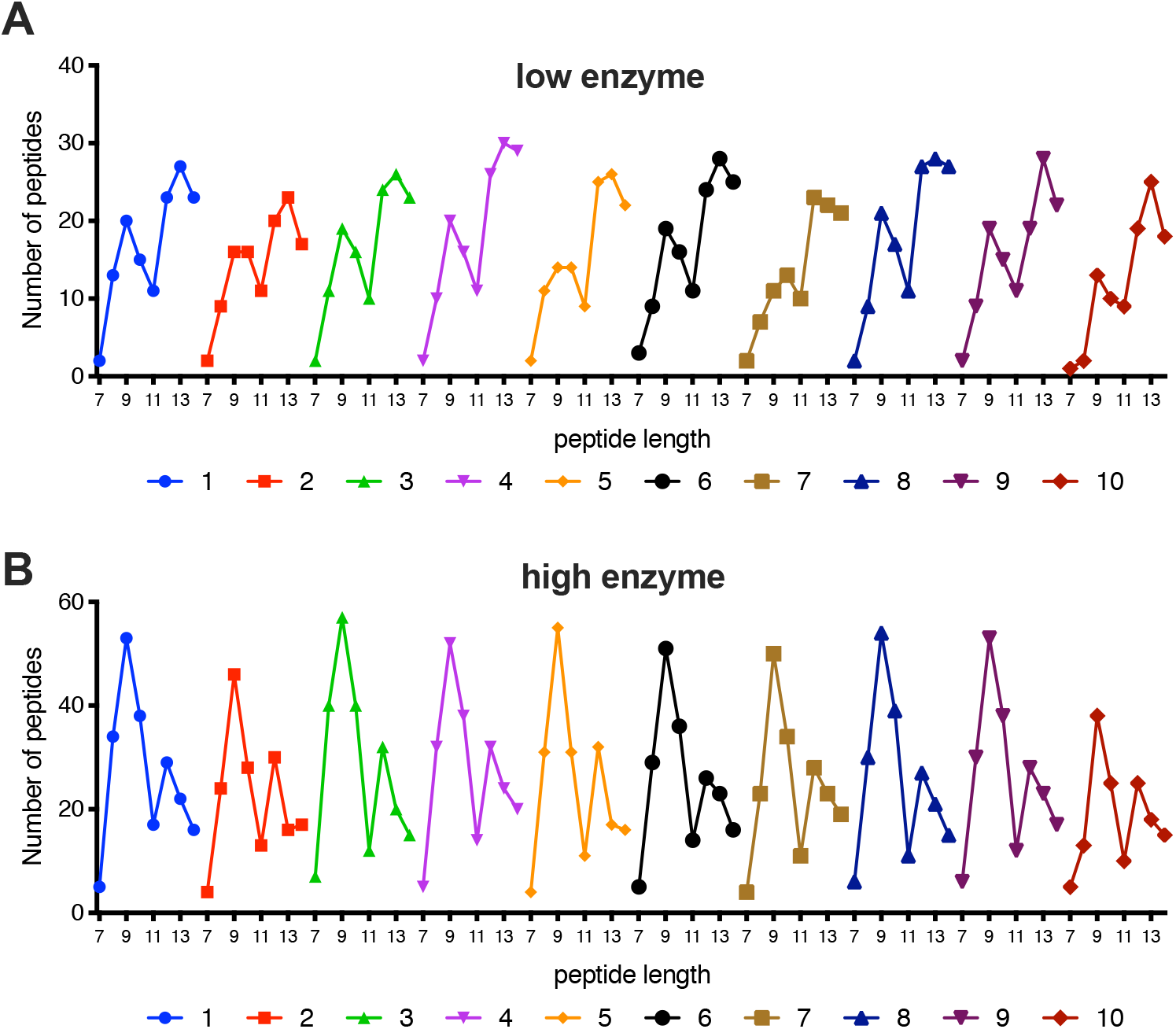
Distribution of peptide length in 7-14mers produced after enzymatic digestion by each of the 10 ERAP1 allotypes, shown for both reaction conditions.

For the low enzyme condition, the distribution of the length of peptides produced by all allotypes formed two apparent “peaks” around 9 and 13 amino acids long respectively (Figure 3A). The peak around 13 amino acids long was much more pronounced suggesting that sequential digestion was still limited for most peptides. Under this condition, allotypes 5, 7 and 10 lagged behind in terms of producing 9mer products.

The overall length distribution was reversed when a higher enzyme concentration was used (Figure 3B). Overall, about double the number of peptides were identified when a higher amount of enzyme was used, consistent with faster sequential digestion of the 15mer substrates. In that condition, the majority of the produced peptides were 9mers for all allotypes, suggesting that all ERAP1 allotypes have the inherent capability to produce peptides with appropriate lengths for MHCI binding. This is consistent with previous reports that have suggested that ERAP1 utilizes a “molecular ruler” mechanism that involves a regulatory allosteric binding site^30,34^. Interestingly, allotypes 5 and 7 that appeared to lag behind in terms of generating 9mers, were now just as effective as the remaining allotypes, suggesting that 9mers can accumulate efficiently for those allotypes, albeit more slowly. In contrast, allotype 10, was again an outlier and produced a lower number of 9mer peptides compared to all the other allotypes.

A key parameter for antigenicity is the sequence of the presented peptides. In this context, it is important to know if the differences between generated peptides are just kinetic or also qualitative (i.e. sequence). To gain insight on whether different ERAP1 allotypes generate different peptide sequences, we calculated for each peptide identified, the number of ERAP1 allotypes that were able to produce it (Figure 4). About double the number of different peptides were produced by the high-enzyme condition, as expected form the enhanced sequential trimming of the 15mer substrates when more enzyme is present. The distribution was found to be U-shaped, in contrast to the expected bell-shaped if peptides were processed in a completely random manner by unrelated protease activities, suggesting specific commonalities and differences in sequence specificity between allotypes. For both reaction conditions, most peptides were produced by all allotypes, indicating that all allotypes have a core sequence preference that shapes the products of the digestion. Strikingly, the second most populous category were peptides that were uniquely produced only by a single allotype (Figure 4A and 4B). This finding suggests that ERAP1 allotypes may carry substrate biases that can influence the repertoire of produced peptides. The generation of allotype-unique peptides suggests that different allotypes can potentially drive different immune responses by generating unique antigenic peptides.

**Figure 4:**
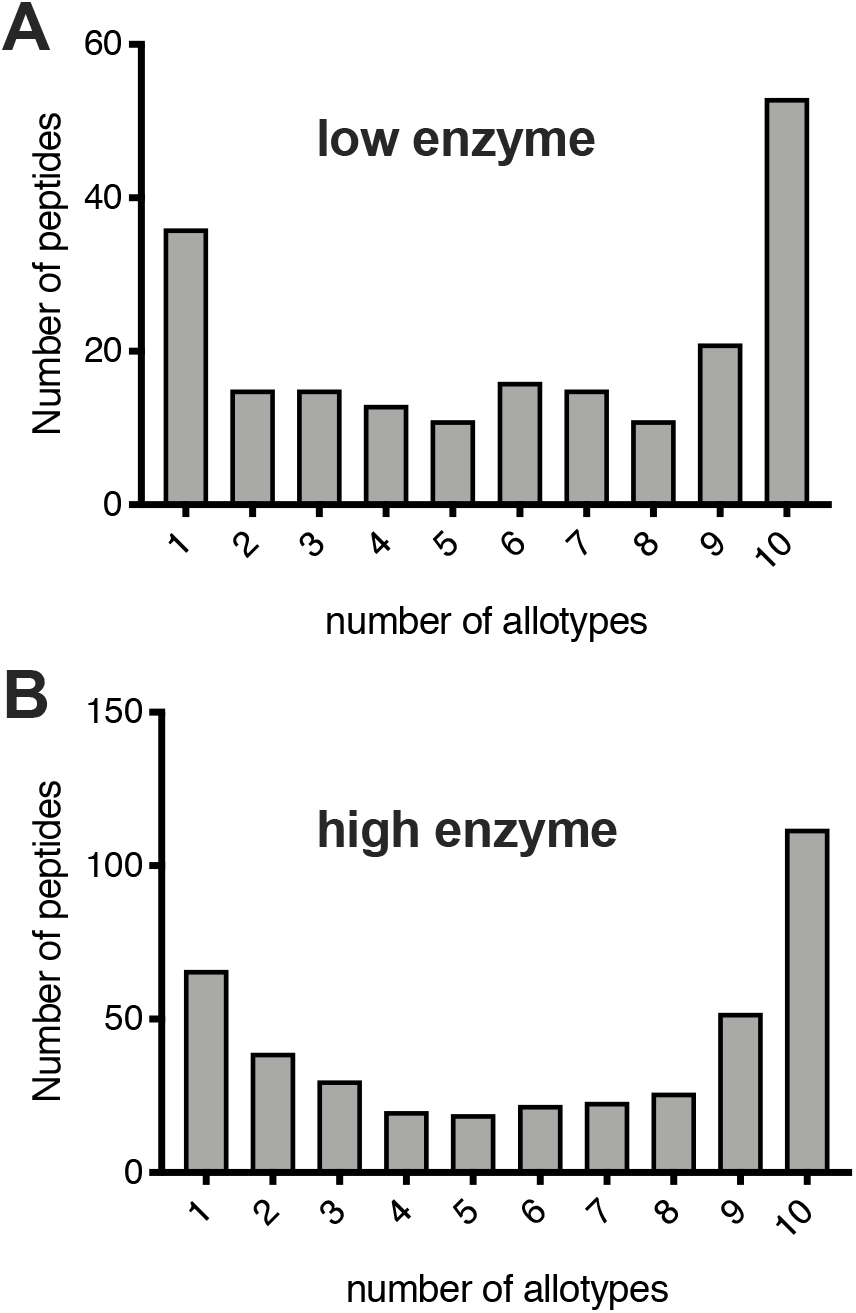
Frequency distribution of produced 7-14mer peptides from both reaction conditions. The number of peptides produced by a specific number of ERAP1 allotypes is shown. For both reaction conditions, the majority of peptides are produced by all 10 allotypes, but a large number of peptides are unique to a specific allotype.

To better understand the commonality of produced peptides for each allotype, we adapted the analysis shown in Figure 4 to report separately for each allotype (Figure 5). For most allotypes and both reaction conditions, the distribution of produced peptides formed a trend towards common peptides, i.e. most of the peptides produced by each allotype were also produced by several others. This was most clear for allotypes 6,8 and 9. Several allotypes however, namely allotypes 1-5, had a secondary peak around peptides that were either unique or produced by 1-3 allotypes in total. A striking outlier was, again, allotype 10. In particular the high enzyme condition, it was the only allotype that the uniqueness distribution was reversed and many of the peptides produced were either unique to allotype 10 or produced by few others. This observation suggests that there could be some sequence bias between allotypes, with allotype 10 being the most extreme example.

**Figure 5:**
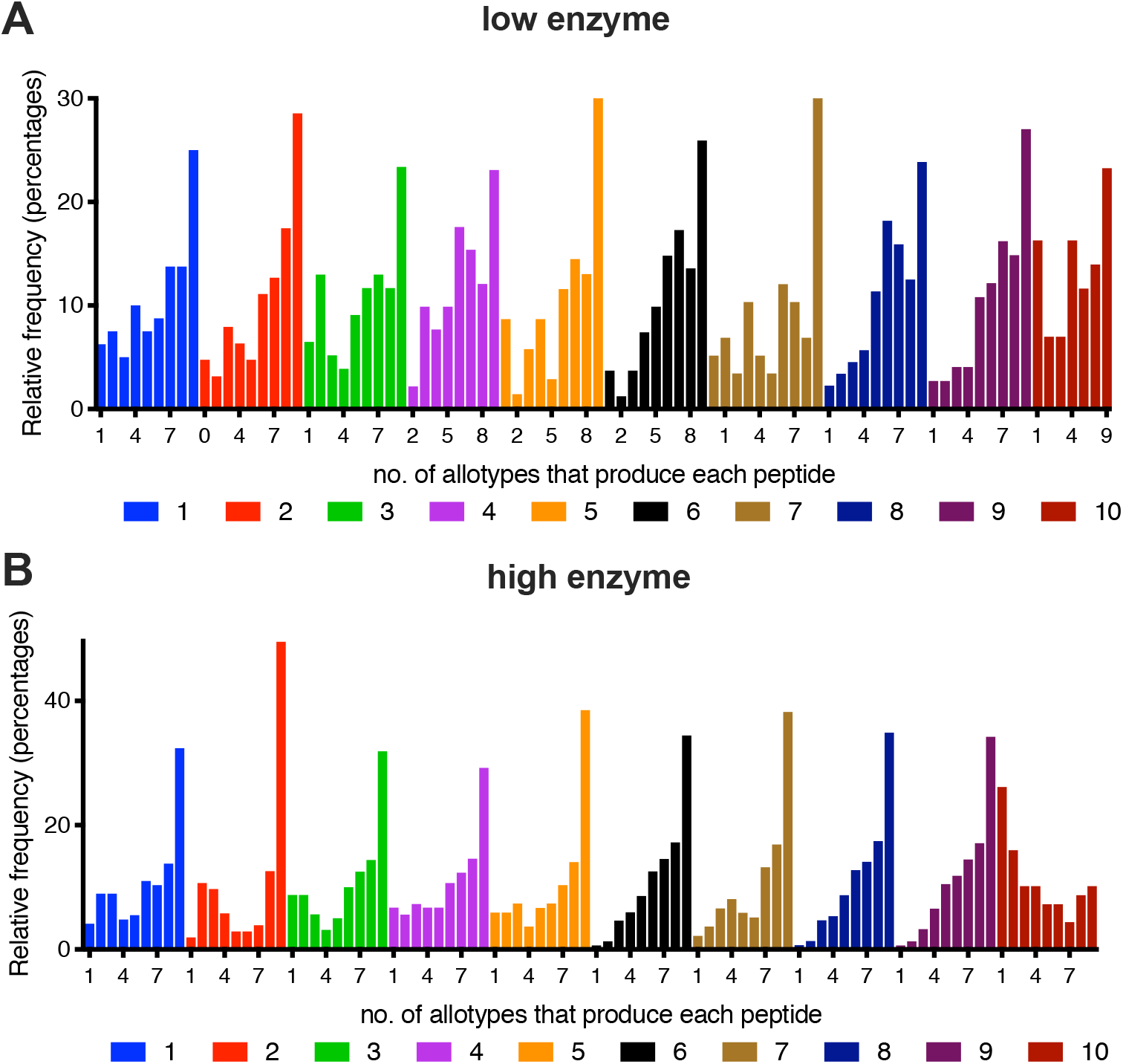
Frequency distribution of generated peptides (7-14mers) produced by one or several different ERAP1 allotypes. The distribution is shown for both reaction conditions and per allotype (1-10). Peptides produced by all 10 allotypes are not plotted.

To further explore the similarities and differences between allotypes in a more unbiased manner we performed principal component analysis (PCA) (Figure 6). With a single exception (one replicate for allotype 9) all replicates clustered together validating the reproducibility of our analyses. For the trimming of the 15mer substrates when using the low enzyme condition, all allotypes and biological replicates cluster together, away from the control reactions (Figure 6A). At the high enzyme condition however, allotype 10 diverged from the other allotypes (Figure 6C). The PCA for the generation of 7-14mers revealed a greater distribution between allotypes. For the low enzyme condition, allotypes 1,3,4,6,8 and 9 formed a cluster. Allotype 10 was a strong outlier, followed by 7 and 2,5. For the high enzyme condition, allotype 10 was again an outlier, followed by 2,5,7 and allotype 3. Overall, PCA revealed that the main difference between allotypes lie on the peptides produced and that allotype 10 consistently appears to differentiate from other allotypes, although intermediate differentiation is observable for some of the other allotypes.

**Figure 6:**
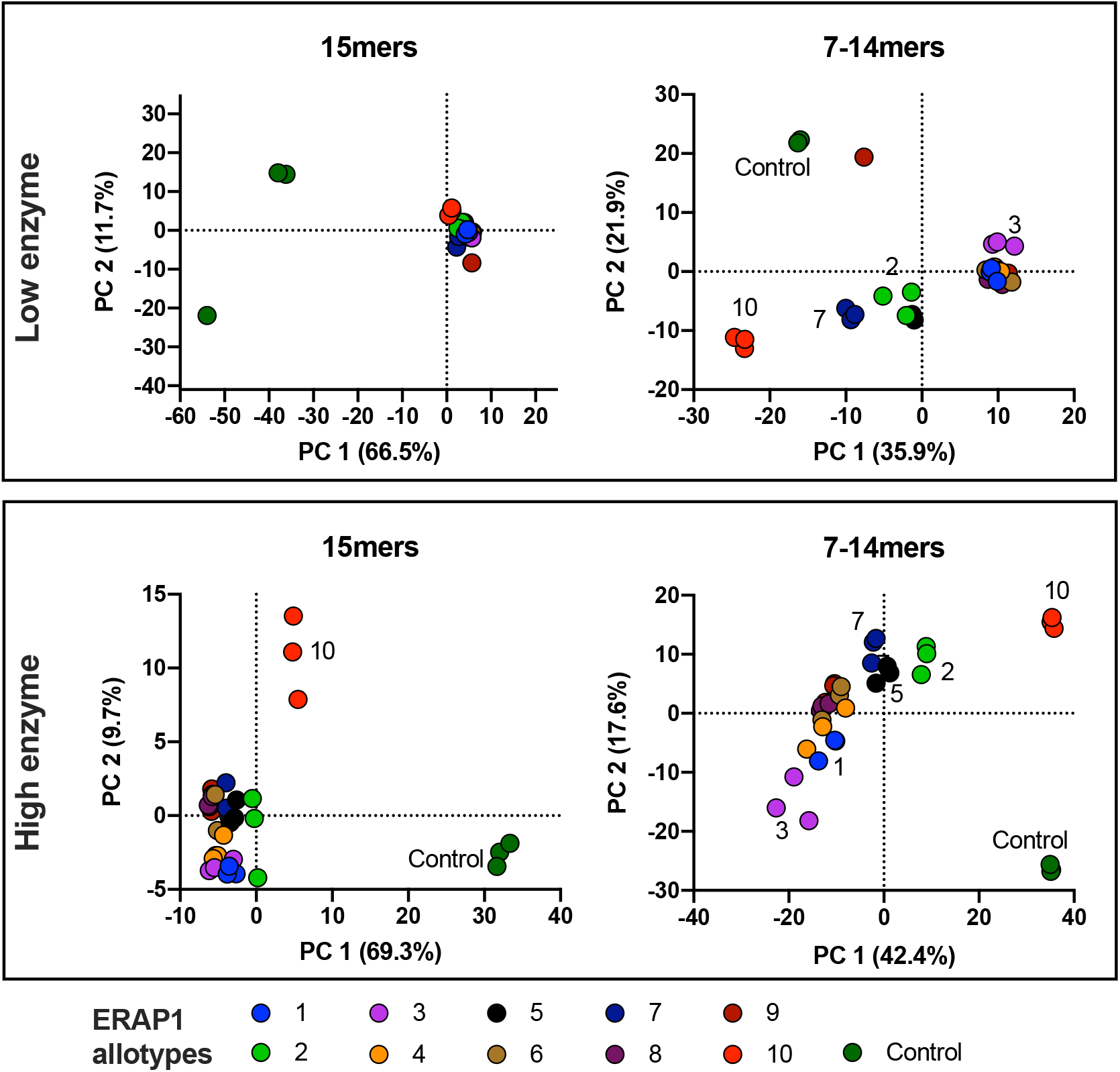
Principal component analysis for the trimming of the initial 15mers and the generation of 7-14mers for both reaction conditions. All biological triplicates (three per allotype) are depicted. The two major components are shown.

The main biological phenomenon that drives cytotoxic T-lymphocyte responses is not intracellular processing but rather antigen presentation by MHC class I molecules (HLA in humans). Which peptides are presented depends on the combination of their availability in the ER and their ability to bind onto the particular HLA-alleles that are expressed in each individual. To evaluate the HLA-binding ability of produced peptides, we utilized a well-established binding prediction server, NetMHCpan-4.1 ^35^. Over 10000 HLA alleles have been discovered to date, having different preferences for binding peptides ^36^. In order to get a reasonable representation of the chances of each generated peptide to be presented, we restricted our analysis to a subset of the most common alleles in the human population, namely HLA-A*01:01, HLA-A*02:01, HLA-A*03:01, HLA-A*24:02, HLA-A*26:01, HLA-B*07:02, HLA-B*08:01, HLA-B*27:05, HLA-B*39:01, HLA-B*40:01, HLA-B*58:01 and HLA-B*15:01. Each of the produced 8-12mers from every ERAP1 allotype was scored for each of the above alleles and the best predicted % rank was plotted for each allotype (Figure 7). A % rank below 2 (cyan region) is considered to be sufficient to promote binding and presentation by the particular allele.

**Figure 7:**
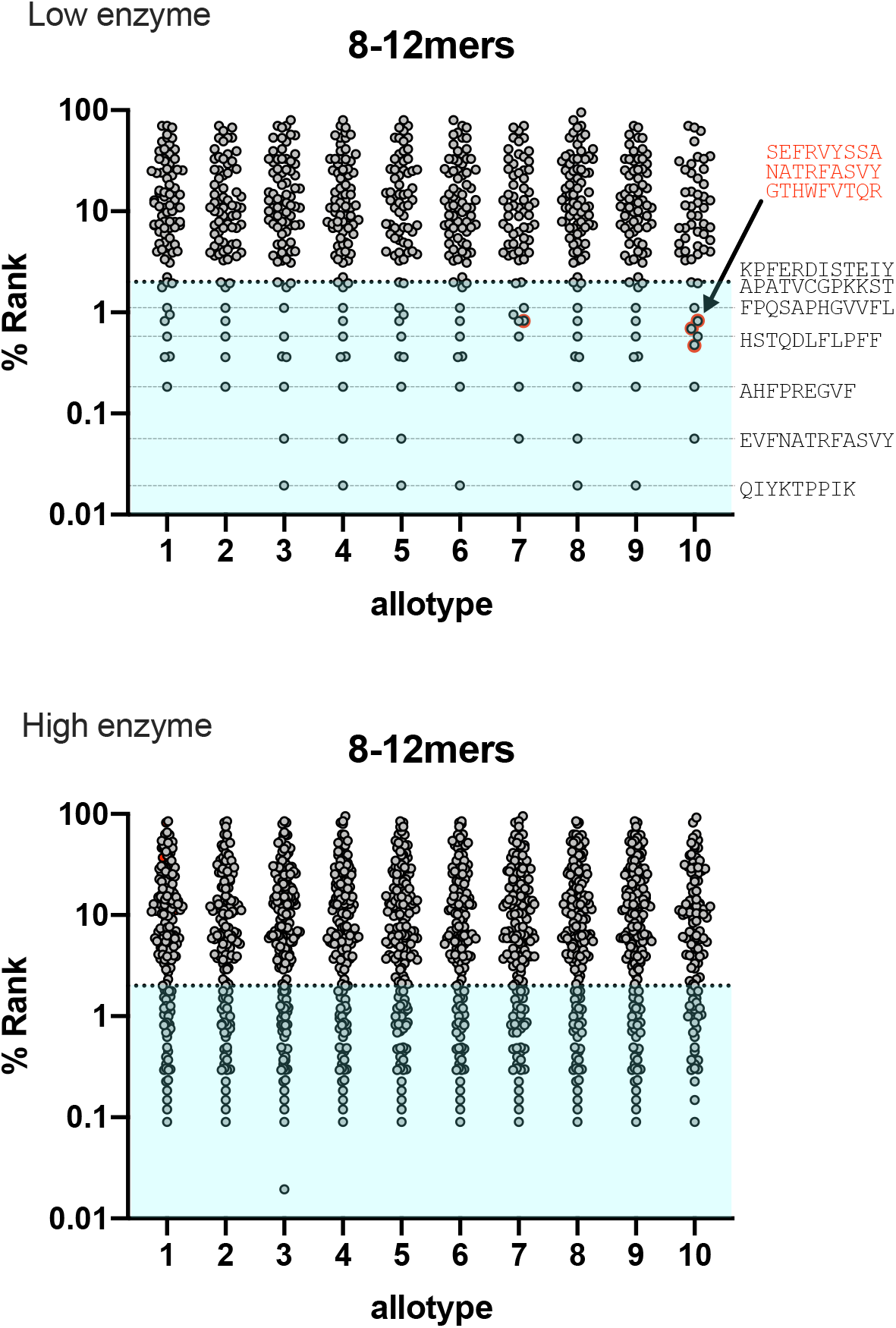
Scatter plots showing the predicted affinity of produced peptides (8-12mers) by both reaction conditions, for common HLA-alleles (HLA-A*01:01, HLA-A*02:01, HLA-A*03:01, HLA-A*24:02, HLA-A*26:01, HLA-B*07:02, HLA-B08:01, HLA-B27:05, HLA-B39:01, HLA-B40:01, HLA-B58:01, HLA-B15:01) as calculated by the NetMHCpan 4.1 Server ^35^.

The majority of produced 8-12mers were not predicted to be binders for the HLA-alleles tested. This however does not mean that they would not be able to bind well onto HLA-alleles not tested here. Not surprisingly, the high enzyme condition produced a much greater number of peptides predicted to bind onto the HLA-alleles tested, possibly because the higher degree of digestion produced more peptides of the appropriate length (such as 9mers, refer to Figure 3B). Overall, all ERAP1 allotypes produced a significant number of peptides that are predicted to bind onto at least one of the HLA-allele tested. This observation included the sub-active allotype 10, which although is less efficient in generating peptides overall, it appears that it is efficient enough to generate many possible candidates for HLA-presentation. Several of the peptides predicted to bind were produced by most or all allotypes (indicated by horizontal dotted lines in Figure 7A), although some peptides were also produced uniquely by a single allotype (indicated by red highlighting in Figure 7A). Some putative HLA-binders produced by the low enzyme condition, were not detected in the high enzyme condition, consistent with the reported ability of ERAP1 to destroy antigenic peptides by over-trimming ^37^. Overall, this analysis shows that although all ERAP1 allotypes are able to generate putative HLA ligands, the exact peptide sequences can be significantly different.

During the last few months intensive research on the adaptive immune responses towards SARS-CoV-2 has allowed the identification of S1 spike glycoprotein epitopes presented by cells of infected individuals^9,38–43^. To correlate the capacity of different ERAP1 allotypes to generate mature epitopes from this antigen, we compared SARS-CoV-2 S1 spike glycoprotein epitopes deposited in the Immune Epitope Database (http://www.iedb.org/) ^36^ with peptides generated during our *in vitro* digestions. Since even transient generation of an epitope could allow for HLA binding, we combined data from both digestions. We identified 20 epitopes that were produced at least from one allotype, 19 of which had published human HLA restrictions (Table 2). Half of those epitopes were generated by all tested ERAP1 allotypes. However, several epitopes were not produced by some allotypes; most notable allotype 10 did not produce 7 of the epitopes. In addition, some epitopes were produced by only a small number of allotypes: epitope SWMESEFRV was produced by allotypes 5 and 10 and epitope NATRFASVY by allotypes 2,7 and 10. Overall, allotype 10 presented a unique fingerprint in terms of its ability to generate known SARS-CoV-2 epitopes, being the least efficient in generating 7 epitopes but at the same time generating 2 epitopes that were not produced by most other ERAP1 allotypes. Overall, our *in vitro* digestions revealed a significant degree of heterogeneity in producing viral antigen epitopes and highlighted a potential unique fingerprint for allotype 10.

**Table 2:**
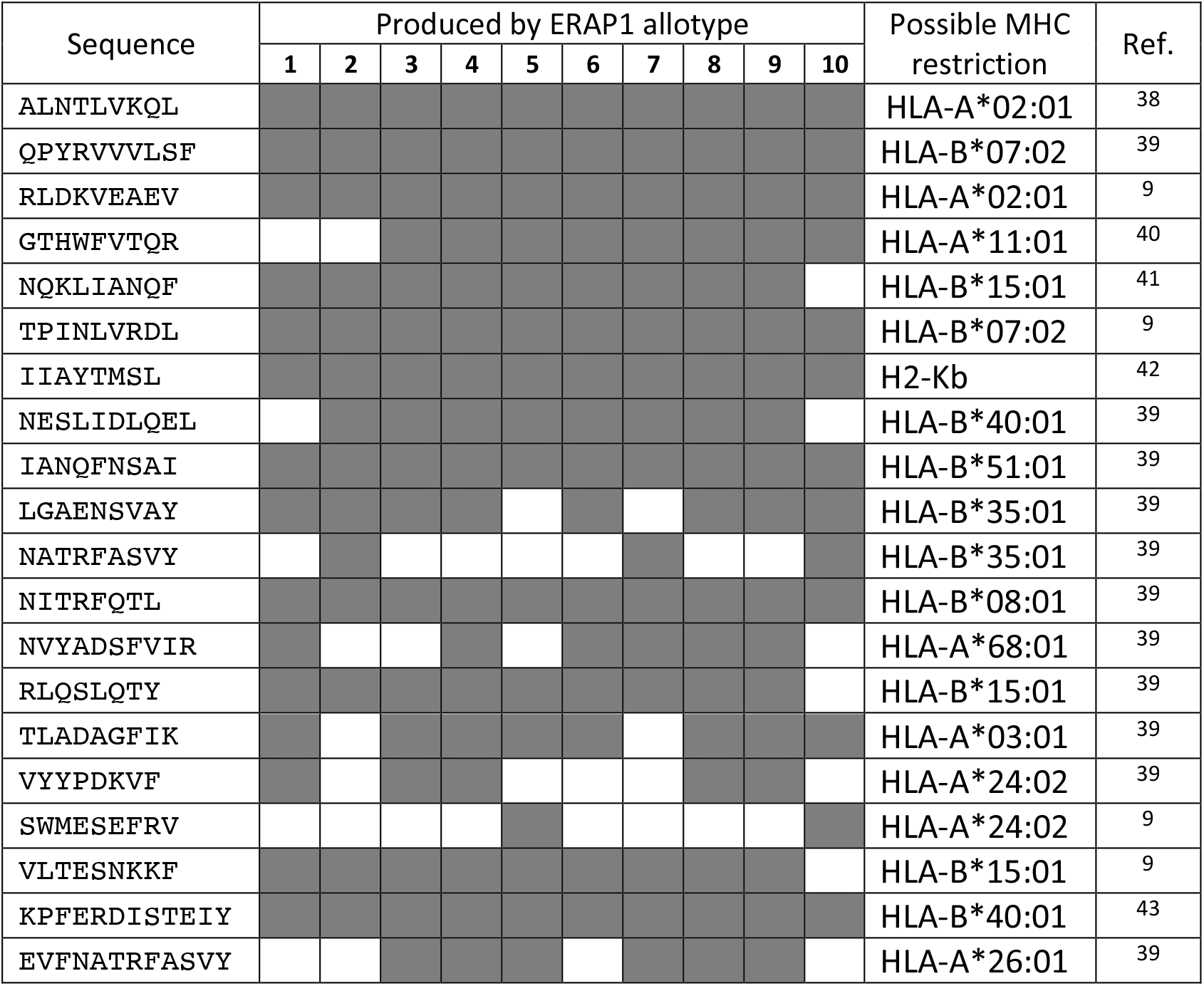
List of known SARD-CoV-2 epitopes identified to be produced by the enzymatic digestion of the S1 glycoprotein peptide pool. The ability of each ERAP1 allotype to produce each of the epitopes is indicated by a greyed box.

## EXPERIMENTAL METHODS

### Materials

The PepMix™ SARS-CoV-2 peptide mixture was purchased by JBT Peptide Technologies GmbH, dissolved in DMSO and stored at -80°C. The two peptide collections (158 and 157 peptides respectively) were mixed at equimolar concentrations and diluted in buffer containing 10 mM Hepes pH 7, 100 mM NaCl to a final concentration of 48 μM.

### Protein Expression and Purification

Recombinant ERAP1 allotypes 1-10 have been described previously ^27^. Briefly, all ERAP1 variants were produced by insect cell culture after infection with recombinant baculovirus and purified to homogeneity by Ni-NTA chromatography and size-exclusion chromatography. Enzymes were stored in aliquots at -80°C with 10% glycerol until needed.

### Enzymatic Reactions

Enzymatic reactions were performed in triplicate in a total volume of 50 μL in 10 mM Hepes pH 7, 150 mM NaCl. Freshly thawed enzyme stocks were added to each reaction to final concentrations of 100 nM or 300 nM (Low enzyme condition and high enzyme condition respectively). Reactions were incubated at 37 °C for 2 h, stopped by the addition of 7.5 μL of a 10% TFA solution, flash frozen in liquid nitrogen, and stored at −80 °C until analyzed by LC-MS/MS.

### LC-MS/MS Analysis

Enzymatic reaction samples were directly injected on a PepSep C18 column (250mm x 0.75mm, 1.9μm) and separated using a gradient of Buffer A (0.1% Formic acid in water) 7% Buffer B (0.1% Formic acid in 80% Acetonitrile) to 35% for 40 min followed by an increase to 45% in 5 min and a second increase to 99% in 0.5min and then kept constant for 4.5min. The column was equilibrated for 15 min prior to the subsequent injection. A full MS was acquired using a Q Exactive HF-X Hybrid Quadropole-Orbitrap mass spectrometer, in the scan range of 350-1500m/z using 120K resolving power with an AGC of 3x 106 and max IT of 100ms, followed by MS/MS scans of the 12 most abundant ions, using 15K resolving power with an AGC of 1x 105 and max IT of 22ms and an NCE of 28 and an dynamic exclusion 0f 30 sec.

### Database Search

The generated raw files were processed by the Proteome Discoverer software (Thermo) (version 2.4) using a workflow for the precursor-based quantification, using SequestHT with Multi Peptide Search and Percolator validation. The Minora algorithm was used for the quantification. The SPIKE_SARS2.fasta was used as database and the search was performed in a unspecific mode eg. No-enzyme specificity was selected. The minimum peptide length was 6 aa. Precursor Mass Tolerance: 10 ppm, Fragment Mass Tolerance: 0.02 Da. Fixed Value PSM Validator was employed for the validation of the peptides and only high confident PSMs were used.

### Statistical Analysis

Three biological replicas of each haplotype were compared against the negative control reaction using the PD pipeline for pairwise ratio protein abundance calculation and the background based t-test.

## DISCUSSION

### Generated peptide repertoire by ERAP1 versus HLA-restricted presentation

It is well established that adaptive immune responses are dependent on cell-surface HLA-restricted antigen presentation. Which peptides are presented depend on two major factors: i) The binding preferences of the HLA alleles expressed by the cell and ii) the availability of peptides with suitable sequence and length. Binding of peptides onto HLA is well-studied, with wany thousands HLA alleles identified to date, having different preferences for peptides ^44^. In addition, several specialized editing chaperones in the ER help select and optimize peptide binding ^45^. In contrast, the mechanisms that control the availability of suitable peptides are less understood and depend on the cellular proteome and the proteolytic cascades that sample it. In particular, proteolytic enzymes like ERAP1 have been shown to influence the immunopeptidome of cells, often in profound manners ^25^. Indeed, ERAP1 SNPs have been shown to both associate with HLA-dependent autoimmunity and cancer ^46^, often in epistasis with HLA, ^47^ and to functionally affect the peptide products ^48,49^. However, functionally-relevant ERAP1 SNPs exist in the population in particular combinations, allotypes, and recent analysis suggested that SNPs can synergize to differentiate allotype functional properties ^27^. Thus, in order to understand and predict antigen presentation, we need to understand the interplay and synergisms between allotypic variation in antigen processing and HLA-restriction.

### Role of ERAP1 allotypic variation in the biology of antigen presentation

The results presented here, clearly suggest that ERAP1 allotypic variation, expressed as 10 common ERAP1 allotypes, translates to the generated repertoire of peptides that can be available for HLA binding. While all allotypes have the capacity to produce a common product “core”, they differentiate in their capacity to produce different sets of peptides that have variable affinities for HLA alleles. In this context, ERAP1 allotypes, parallel the capacity of HLA for defining the immunopeptidome in qualitative but not absolute patterns and likely complement and synergize with HLA haplotypes to shape the antigen presentation “preferences” of each cell. In this context, a more fundamental biological role for ERAP1 can emerge. While initial studies on ERAP1 highlighted its importance for the generation of particular epitopes ^50–52^, its role in destroying some antigenic epitopes later emerged as equally important ^53,54^. However, more global immunopeptidomic analyses, later demonstrated that the effect or ERAP1 can be limited ^37,55,56^ or more focused on antigenic peptide destruction ^37,57^. Thus, it is possible that the main biological function of ERAP1 is not to be an obligate generator of antigenic peptides as initially suspected, but rather lies in its allotypic variability. Synergism between ERAP1 trimming specificity and HLA binding preferences could enhance the variability of HLA-presentation, thus contributing to the variability of adaptive immune responses in the human population. In this context, the term “ERAP1-dependend and HLA-restricted” may be a more appropriate way to describe the breadth of antigen presentation for a given cell or individual that carry a limited set of ERAP1 allotypes and HLA-haplotypes.

### Allotype 10 is not a loss-of-function allele but produces a different fingerprint of peptides

In a previous study, we demonstrated that allotype 10 had a significantly lower enzymatic activity for many substrates tested and this was due to both reduction in catalytic efficiency as well as substrate affinity ^27^. That study however was performed with a limited number of substates analyzed one at a time. Since however ERAP1-substrate interactions can be complex and include competition between substrates and phenomena such as substrate inhibition^31^, concurrent analysis of digestion of multiple peptides can provide more accurate insight and better emulate the sub-cellular environment where ERAP1 normally functions. Indeed, when analyzing the concurrent digestion of hundreds of substrates, allotype 10 was found to be only about 2-fold less enzymatically efficient. Still, allotype 10 was found to produce peptides with distinctly different patterns and constitute a functional outlier compared to the other allotypes, suggesting mechanistical differences in substrate selection. Thus, the notion that individuals homozygous for allotype 10, carry just a sub-active ERAP1 enzyme, should be revised since it appears to only apply to particular substrates. Rather, homozygous individuals appear to carry an ERAP1 variant that exerts different specificity pressures on the peptide repertoire and could result to particular changes in presented peptides.

The structural basis of the distinct behavior of allotype 10 is not clear. Some insight however can be derived by comparing the SNPs present in this allotype to the SNPs in other allotypes. In particular, allotype 10 contains a combination of two SNPs that are unique for this allotype, namely Val349 and Gln725. Both of those residues were found to either directly or indirectly interact with substrate analogues in recently determined crystal structures of ERAP1 ^30^. Val349 is located adjacent to the active site, and although a conservative substitution (Met to Val) could influence substrate recognition. Gln725 is a non-conservative substitution (Arg to Gln) and lies near the hinge domain of ERAP1 and adjacent to the C-terminal residue of a 10mer peptide substrate analogue that has been crystallized with ERAP1 ^30^. The SNP Arg725 makes a salt-bridge with Asp766 which interacts with the C-terminal Lys residue of that peptide. Furthermore Gln725 lies in a region that undergoes significant structural reconfigurations when ERAP1 changes conformation from the “open” state to its “close” state and could thus influence turnover rates ^58^. It is therefore possible that the combination of those two SNPs in allotype 10, synergizes to affect catalytic rates for different substrates, resulting in an apparent different specificity. Additional structural studies will be necessary to better understand the unique behavior of this ERAP1 allotype.

### ERAP1 allotypic variation and COVID-19

The significant variability of individuals to susceptibility to COVID-19 has been a hallmark of the current pandemic^10^. While age appears to be the most important factor, the genetic blueprint of infected individuals also appears to predispose some to severe COVID-19, but the exact genetic factors responsible are poorly understood. Studies have shown that rare mutations as well as polymorphic variability in immune system components can be critical. The role of rare mutations in components of the innate immune response and in particular the interferon I pathway ^13,14^, has been described but the exact role of natural polymorphic variation in components of adaptive immunity, such as HLA-alleles, in only now emerging, possibly due to difficulties in isolating specific genetic associations from complex polymorphic genetic systems by population studies ^15–17^. The existence and potential role of ERAP1 allotypes have only recently been recognized ^26^ and thus have not been studied in the context of COVID-19 susceptibility. Our results, however, suggest that ERAP1 allotypic variation could play, in tandem to HLA-alleles, important roles in determining anti-SARS-CoV-2 immune responses and by extension, susceptibility to severe COVID-19. For example, allotype 10 homozygous individuals, which could represent up to 4.8% of the population in Europe ^27^, could be defective in producing epitopes from the S1 glycoprotein that are presented from particular HLA alleles. Conversely, our analysis suggests that particular antigenic epitopes may be produced by only specific ERAP1 allotypes. Thus, particular combinations of ERAP1 allotypes with HLA alleles could lead to either effective or defective adaptive immune responses against SARS-CoV-2. It is therefore likely important to include the ERAP1 allotypic state in genetic analyses that focus on HLA-associations with predisposition to severe COVID-19.

### Limitations of the study

While our study highlights the potential importance of ERAP1 allotypes in immune response variability between individuals, it has some important limitations that need to be taken into account when interpreting results. The *in vitro* nature of the digestions may not be an optimal surrogate for cellular antigen processing, either due to missing components (such as other peptidases) or due to cellular compartmentalization. HLA-restriction is only indirectly simulated and thus does not take into account kinetic components of peptide competition for binding inside the ER. Furthermore, although antigenic peptide destruction is evident when comparing the two reaction conditions, it is difficult to analyze statistically without an explicit set of 9mer peptides in the initial reaction. Finally, although our approach of concurrently analyzing hundreds of peptide-trimming reactions proved invaluable in identifying difference between trimming patterns of ERAP1 allotypes, the number of peptides used was not sufficient to allow for the explicit identification of sequence motifs in trimming preferences. The latter however may not be readily feasible given the large substrate cavity of ERAP1 that allows for a complex landscape of peptide-enzyme interactions^30^.

### Cytotoxic responses after vaccination – different response depending on ERAP1 allotype?

Although vaccines developed for COVID-19 had as a primary goal the induction of robust antibody responses, the generation of potent and long-lasting cellular adaptive responses will be critical for controlling disease severity and long-term immunity ^59–61^.

Although not yet established, some peptide epitopes from SARS-CoV-2 antigens may be immunodominant and shape long-term immunity for a large percentage of individuals. Our results here suggest that the polymorphic variability in peptide epitope generation by ERAP1 allotypes could also play a role, in tandem to HLA-restriction, in eliciting and sustaining vaccine-induced cellular immunity. Further studies aiming at defining SARS-CoV-2 immunodominant epitopes shared across individuals will be necessary to test this hypothesis.

## Conclusions

We demonstrate that ERAP1 allotypes common in the population, demonstrate significant differences in their ability to process antigenic epitopes precursors derived from the S1 spike glycoprotein of SARS-CoV-2. A clear outlier is allotype 10, a common ERAP1 variant (present in more than 20% of Europeans) previously suggested to be sub-active, which we find to still be able to generate HLA ligands but with distinct patterns. Our findings suggest that ERAP1 allotypes can be a major contributor to heterogeneity in antigen presentation and in conjunction with HLA-allele binding specificity, contribute to variable immune responses to disease including COVID-19.

## ACKNOWLEDGMENTS

The authors would like to extend their gratitude to GlaxoSmithKline and in particular Drs. Semra Kitchen and Jonathan P. Hutchinson for providing recombinant enzymes that allowed for this study to be performed in a timely manner^27^.

## AUTHOR CONTRIBUTIONS

G.S., M.S. and G.P. performed LC-MS/MS analysis, analyzed data and interpreted data. I.T. performed enzymatic digestions and interpreted data. E.S. performed experiments, analyzed and interpreted data and wrote the paper with help from all authors. All authors have approved the final version of the manuscript.

## DATA DEPOSITION

The mass spectrometry proteomics data have been deposited to the ProteomeXchange Consortium via the PRIDE^62^ partner repository with the data set identifier PXD027006 (http://www.ebi.ac.uk/pride/archive/).

## REFERENCES

(1) Brodin, P. Immune Determinants of COVID-19 Disease Presentation and Severity. Nature Medicine. Nature Research January 1, 2021, pp 28–33. https://doi.org/10.1038/s41591-020-01202-8.

(2) St John, A. L.; Rathore, A. P. S. Early Insights into Immune Responses during COVID-19. J. Immunol. 2020, 205 (3), 555–564. https://doi.org/10.4049/jimmunol.2000526.

(3) Song, P.; Li, W.; Xie, J.; Hou, Y.; You, C. Cytokine Storm Induced by SARS-CoV-2. Clin. Chim. Acta 2020, 509, 280–287. https://doi.org/10.1016/j.cca.2020.06.017.

(4) Li, K.; Hao, Z.; Zhao, X.; Du, J.; Zhou, Y. SARS-CoV-2 Infection-Induced Immune Responses: Friends or Foes? Scand. J. Immunol. 2020, 92 (2), e12895. https://doi.org/10.1111/sji.12895.

(5) Radbruch, A.; Chang, H.-D. A Long-Term Perspective on Immunity to COVID. Nature 2021. https://doi.org/10.1038/d41586-021-01557-z.

(6) Leslie, M. T Cells Found in Coronavirus Patients “bode Well” for Long-Term Immunity. Science (80-.). 2020, 368 (6493), 809–810. https://doi.org/10.1126/science.368.6493.809.

(7) Tarke, A.; Sidney, J.; Kidd, C. K.; Dan, J. M.; Ramirez, S. I.; Yu, E. D.; Mateus, J.; da Silva Antunes, R.; Moore, E.; Rubiro, P.; Methot, N.; Phillips, E.; Mallal, S.; Frazier, A.; Rawlings, S. A.; Greenbaum, J. A.; Peters, B.; Smith, D. M.; Crotty, S.; Weiskopf, D.; Grifoni, A.; Sette, A. Comprehensive Analysis of T Cell Immunodominance and Immunoprevalence of SARS-CoV-2 Epitopes in COVID-19 Cases. Cell Reports Med. 2021, 2 (2). https://doi.org/10.1016/j.xcrm.2021.100204.

(8) Ledford, H. How “killer” T Cells Could Boost COVID Immunity in Face of New Variants. Nature. NLM (Medline) February 1, 2021, pp 374–375. https://doi.org/10.1038/d41586-021-00367-7.

(9) Saini, S. K.; Hersby, D. S.; Tamhane, T.; Povlsen, H. R.; Amaya Hernandez, S. P.; Nielsen, M.; Gang, A. O.; Hadrup, S. R. SARS-CoV-2 Genome-Wide T Cell Epitope Mapping Reveals Immunodominance and Substantial CD8+ T Cell Activation in COVID-19 Patients. Sci. Immunol. 2021, 6 (58). https://doi.org/10.1126/sciimmunol.abf7550.

(10) Pereira, N. L.; Ahmad, F.; Byku, M.; Cummins, N. W.; Morris, A. A.; Owens, A.; Tuteja, S.; Cresci, S. COVID-19: Understanding Inter-Individual Variability and Implications for Precision Medicine. Mayo Clinic Proceedings. Elsevier Ltd February 1, 2021, pp 446– 463. https://doi.org/10.1016/j.mayocp.2020.11.024.

(11) Fricke-Galindo, I.; Falfán-Valencia, R. Genetics Insight for COVID-19 Susceptibility and Severity: A Review. Frontiers in Immunology. Frontiers Media S.A. April 1, 2021, p 1057. https://doi.org/10.3389/fimmu.2021.622176.

(12) Zeberg, H.; Pääbo, S. The Major Genetic Risk Factor for Severe COVID-19 Is Inherited from Neanderthals. Nature 2020, 587 (7835), 610–612. https://doi.org/10.1038/s41586-020-2818-3.

(13) Zhang, Q.; Liu, Z.; Moncada-Velez, M.; Chen, J.; Ogishi, M.; Bigio, B.; Yang, R.; Arias, A. A.; Zhou, Q.; Han, J. E.; Ugurbil, A. C.; Zhang, P.; Rapaport, F.; Zhang, X.; et al. Inborn Errors of Type I IFN Immunity in Patients with Life-Threatening COVID-19. Science (80-.). 2020, 370 (6515). https://doi.org/10.1126/science.abd4570.

(14) Lee, J. S.; Shin, E. C. The Type I Interferon Response in COVID-19: Implications for Treatment. Nature Reviews Immunology. Nature Research October 1, 2020, pp 585– 586. https://doi.org/10.1038/s41577-020-00429-3.

(15) Shkurnikov, M.; Nersisyan, S.; Jankevic, T.; Galatenko, A.; Gordeev, I.; Vechorko, V.; Tonevitsky, A. Association of HLA Class I Genotypes With Severity of Coronavirus Disease-19. Front. Immunol. 2021, 12, 641900. https://doi.org/10.3389/fimmu.2021.641900.

(16) Saulle, I.; Vicentini, C.; Clerici, M.; Biasin, M. Antigen Presentation in SARS-CoV-2 Infection: The Role of Class I HLA and ERAP Polymorphisms. Hum. Immunol. 2021. https://doi.org/10.1016/j.humimm.2021.05.003.

(17) Secolin, R.; de Araujo, T. K.; Gonsales, M. C.; Rocha, C. S.; Naslavsky, M.; Marco, L. De; Bicalho, M. A. C.; Vazquez, V. L.; Zatz, M.; Silva, W. A.; Lopes-Cendes, I. Genetic Variability in COVID-19-Related Genes in the Brazilian Population. Hum. Genome Var. 2021, 8 (1), 15. https://doi.org/10.1038/s41439-021-00146-w.

(18) Rock, K. L.; Goldberg, A. L. Degradation of Cell Proteins and the Generation of MHC Class I-Presented Peptides. Annu Rev Immunol 1999, 17, 739–779.

(19) Weimershaus, M.; Evnouchidou, I.; Saveanu, L.; van Endert, P. Peptidases Trimming MHC Class I Ligands. Curr. Opin. Immunol. 2013, 25 (1), 90–96. https://doi.org/10.1016/j.coi.2012.10.001.

(20) Hammer, G. E.; Kanaseki, T.; Shastri, N. The Final Touches Make Perfect the Peptide-MHC Class I Repertoire. Immunity 2007, 26 (4), 397–406. https://doi.org/10.1016/j.immuni.2007.04.003.

(21) Reeves, E.; Wood, O.; Ottensmeier, C. H.; King, E. V; Thomas, G. J.; Elliott, T.; James, E. HPV Epitope Processing Differences Correlate with ERAP1 Allotype and Extent of CD8(+) T-Cell Tumor Infiltration in OPSCC. Cancer Immunol Res 2019, 7 (7), 1202–1213. https://doi.org/10.1158/2326-6066.CIR-18-0498.

(22) Kim, S.; Lee, S.; Shin, J.; Kim, Y.; Evnouchidou, I.; Kim, D.; Kim, Y.-K.; Kim, Y.-E.; Ahn, J.-H.; Riddell, S. R.; Stratikos, E.; Kim, V. N.; Ahn, K. Human Cytomegalovirus MicroRNA MiR-US4-1 Inhibits CD8 ^+^ T Cell Responses by Targeting the Aminopeptidase ERAP1. Nat. Immunol. 2011, 12 (10). https://doi.org/10.1038/ni.2097.

(23) Robinson, J.; Barker, D. J.; Georgiou, X.; Cooper, M. A.; Flicek, P.; Marsh, S. G. E. IPD-IMGT/HLA Database. Nucleic Acids Res. 2020, 48 (D1), D948–D955. https://doi.org/10.1093/nar/gkz950.

(24) Kirino, Y.; Bertsias, G.; Ishigatsubo, Y.; Mizuki, N.; Tugal-Tutkun, I.; Seyahi, E.; Ozyazgan, Y.; Sacli, F. S.; Erer, B.; Inoko, H.; Emrence, Z.; Cakar, A.; Abaci, N.; Ustek, D.; Satorius, C.; Ueda, A.; Takeno, M.; Kim, Y.; Wood, G. M.; Ombrello, M. J.; Meguro, A.; Gul, A.; Remmers, E. F.; Kastner, D. L. Genome-Wide Association Analysis Identifies New Susceptibility Loci for Behcet’s Disease and Epistasis between HLA-B*51 and ERAP1. Nat Genet 2013, 45 (2), 202–207. https://doi.org/10.1038/ng.2520ng.2520 [pii].

(25) López de Castro, J.A. How ERAP1 and ERAP2 Shape the Peptidomes of Disease-Associated MHC-I Proteins. Front. Immunol. 2018, 9 (October), 2463. https://doi.org/10.3389/fimmu.2018.02463.

(26) Ombrello, M. J.; Kastner, D. L.; Remmers, E. F. Endoplasmic Reticulum-Associated Amino-Peptidase 1 and Rheumatic Disease: Genetics. Curr. Opin. Rheumatol. 2015, 27 (4), 349–356. https://doi.org/10.1097/BOR.0000000000000189.

(27) Hutchinson, J. P.; Temponeras, I.; Kuiper, J.; Cortes, A.; Korczynska, J.; Kitchen, S.; Stratikos, E. Common Allotypes of ER Aminopeptidase 1 Have Substrate-Dependent and Highly Variable Enzymatic Properties. J. Biol. Chem. 2021, 296, 100443. https://doi.org/10.1016/j.jbc.2021.100443.

(28) Reeves, E.; Edwards, C. J.; Elliott, T.; James, E. Naturally Occurring ERAP1 Haplotypes Encode Functionally Distinct Alleles with Fine Substrate Specificity. J Immunol 2013, 191 (1), 35–43. https://doi.org/10.4049/jimmunol.1300598jimmunol.1300598[pii].

(29) Roberts, A. R.; Appleton, L. H.; Cortes, A.; Vecellio, M.; Lau, J.; Watts, L.; Brown, M. A.; Wordsworth, P. ERAP1 Association with Ankylosing Spondylitis Is Attributable to Common Genotypes Rather than Rare Haplotype Combinations. Proc Natl Acad Sci U S A 2017, 114 (3), 558–561. https://doi.org/10.1073/pnas.1618856114.

(30) Giastas, P.; Mpakali, A.; Papakyriakou, A.; Lelis, A.; Kokkala, P.; Neu, M.; Rowland, P.; Liddle, J.; Georgiadis, D.; Stratikos, E. Mechanism for Antigenic Peptide Selection by Endoplasmic Reticulum Aminopeptidase 1. Proc. Natl. Acad. Sci. U. S. A. 2019, 116 (52). https://doi.org/10.1073/pnas.1912070116.

(31) Evnouchidou, I.; Kamal, R. P.; Seregin, S. S.; Goto, Y.; Tsujimoto, M.; Hattori, A.; Voulgari, P. V; Drosos, A. A.; Amalfitano, A.; York, I. A.; Stratikos, E. Coding Single Nucleotide Polymorphisms of Endoplasmic Reticulum Aminopeptidase 1 Can Affect Antigenic Peptide Generation in Vitro by Influencing Basic Enzymatic Properties of the Enzyme. J Immunol 2011, 186 (4), 1909–1913. https://doi.org/jimmunol.1003337[pii]10.4049/jimmunol.1003337.

(32) Evnouchidou, I.; Kamal, R. P.; Seregin, S. S.; Goto, Y.; Tsujimoto, M.; Hattori, A.; Voulgari, P. V.; Drosos, A. A.; Amalfitano, A.; York, I. A.; Stratikos, E. Cutting Edge: Coding Single Nucleotide Polymorphisms of Endoplasmic Reticulum Aminopeptidase 1 Can Affect Antigenic Peptide Generation in Vitro by Influencing Basic Enzymatic Properties of the Enzyme. J. Immunol. 2011, 186 (4). https://doi.org/10.4049/jimmunol.1003337.

(33) Stamatakis, G.; Samiotaki, M.; Mpakali, A.; Panayotou, G.; Stratikos, E. Generation of SARS-CoV-2 S1 Spike Glycoprotein Putative Antigenic Epitopes in Vitro by Intracellular Aminopeptidases. J. Proteome Res. 2020, 19 (11), 4398–4406. https://doi.org/10.1021/acs.jproteome.0c00457.

(34) Chang, S. C.; Momburg, F.; Bhutani, N.; Goldberg, A. L. The ER Aminopeptidase, ERAP1, Trims Precursors to Lengths of MHC Class I Peptides by a “Molecular Ruler” Mechanism. Proc Natl Acad Sci U S A 2005, 102 (47), 17107–17112.

(35) Reynisson, B.; Alvarez, B.; Paul, S.; Peters, B.; Nielsen, M. NetMHCpan-4.1 and NetMHCIIpan-4.0: Improved Predictions of MHC Antigen Presentation by Concurrent Motif Deconvolution and Integration of MS MHC Eluted Ligand Data. Nucleic Acids Res. 2020, 48 (W1), W449–W454. https://doi.org/10.1093/nar/gkaa379.

(36) Vita, R.; Mahajan, S.; Overton, J. A.; Dhanda, S. K.; Martini, S.; Cantrell, J. R.; Wheeler, D. K.; Sette, A.; Peters, B. The Immune Epitope Database (IEDB): 2018 Update. Nucleic Acids Res. 2019, 47 (D1), D339–D343. https://doi.org/10.1093/nar/gky1006.

(37) Admon, A. ERAP1 Shapes Just Part of the Immunopeptidome. Hum. Immunol. 2019, 80 (5), 296–301. https://doi.org/10.1016/j.humimm.2019.03.004.

(38) Shomuradova, A. S.; Vagida, M. S.; Sheetikov, S. A.; Zornikova, K. V.; Kiryukhin, D.; Titov, A.; Peshkova, I. O.; Khmelevskaya, A.; Dianov, D. V.; Malasheva, M.; Shmelev, A.; Serdyuk, Y.; Bagaev, D. V.; Pivnyuk, A.; Shcherbinin, D. S.; Maleeva, A. V.; Shakirova, N. T.; Pilunov, A.; Malko, D. B.; Khamaganova, E. G.; Biderman, B.; Ivanov, A.; Shugay, M.; Efimov, G. A. SARS-CoV-2 Epitopes Are Recognized by a Public and Diverse Repertoire of Human T Cell Receptors. Immunity 2020, 53 (6), 1245-1257.e5. https://doi.org/10.1016/j.immuni.2020.11.004.

(39) Tarke, A.; Sidney, J.; Kidd, C. K.; Dan, J. M.; Ramirez, S. I.; Yu, E. D.; Mateus, J.; da Silva Antunes; R., Moore, E.; Rubiro, P.; Methot, N.; Phillips, E.; Mallal, S.; Frazier, A.; Rawlings, S. A.; Greenbaum, J. A.; Peters, B.; Smith, D. M.; Crotty, S.; Weiskopf, D.; Grifoni, A.; Sette, A. Comprehensive Analysis of T Cell Immunodominance and Immunoprevalence of SARS-CoV-2 Epitopes in COVID-19 Cases. Cell reports. Med. 2021, 2 (2), 100204. https://doi.org/10.1016/j.xcrm.2021.100204.

(40) Kared, H.; Redd, A. D.; Bloch, E. M.; Bonny, T. S.; Sumatoh, H.; Kairi, F.; Carbajo, D.; Abel, B.; Newell, E. W.; Bettinotti, M. P.; Benner, S. E.; Patel, E. U.; Littlefield, K.; Laeyendecker, O.; Shoham, S.; Sullivan, D.; Casadevall, A.; Pekosz, A.; Nardin, A.; Fehlings, M.; Tobian, A. A. R.; Quinn, T. C. SARS-CoV-2-Specific CD8+ T Cell Responses in Convalescent COVID-19 Individuals. J. Clin. Invest. 2021, 131 (5). https://doi.org/10.1172/JCI145476.

(41) Minervina, A. A.; Komech, E. A.; Titov, A.; Koraichi, M. B.; Rosati, E.; Mamedov, I. Z.; Franke, A.; Efimov, G. A.; Chudakov, D. M.; Mora, T.; Walczak, A. M.; Lebedev, Y. B.; Pogorelyy, M. V. Longitudinal High-Throughput Tcr Repertoire Profiling Reveals the Dynamics of t-Cell Memory Formation after Mild Covid-19 Infection. Elife 2021, 10, 1– 17. https://doi.org/10.7554/eLife.63502.

(42) Hassert, M.; Geerling, E.; Stone, E. T.; Steffen, T. L.; Feldman, M. S.; Dickson, A. L.; Class, J.; Richner, J. M.; Brien, J. D.; Pinto, A. K. MRNA Induced Expression of Human Angiotensin-Converting Enzyme 2 in Mice for the Study of the Adaptive Immune Response to Severe Acute Respiratory Syndrome Coronavirus 2. PLoS Pathog. 2020, 16 (12). https://doi.org/10.1371/journal.ppat.1009163.

(43) Snyder, T. M.; Gittelman, R. M.; Klinger, M.; May, D. H.; Osborne, E. J.; Taniguchi, R.; Zahid, H. J.; Kaplan, I. M.; Dines, J. N.; Noakes, M. N.; Pandya, R.; Chen, X.; Elasady, S.; Robins, H. S.; et al. Magnitude and Dynamics of the T-Cell Response to SARS-CoV-2 Infection at Both Individual and Population Levels. medRxiv Prepr. Serv. Heal. Sci. 2020. https://doi.org/10.1101/2020.07.31.20165647.

(44) Robinson, J.; Halliwell, J. A.; Hayhurst, J. D.; Flicek, P.; Parham, P.; Marsh, S. G. E. The IPD and IMGT/HLA Database: Allele Variant Databases. Nucleic Acids Res. 2015, 43 (D1), D423–D431. https://doi.org/10.1093/nar/gku1161.

(45) Thomas, C.; Tampe, R. MHC I Chaperone Complexes Shaping Immunity. Curr. Opin. Immunol. 2019, 58, 9–15. https://doi.org/10.1016/j.coi.2019.01.001.

(46) Reeves, E.; Islam, Y.; James, E. ERAP1: A Potential Therapeutic Target for a Myriad of Diseases. Expert Opin Ther Targets 2020, 24 (6), 535–544. https://doi.org/10.1080/14728222.2020.1751821.

(47) Cortes, A.; Pulit, S. L.; Leo, P. J.; Pointon, J. J.; Robinson, P. C.; Weisman, M. H.; Ward, M.; Gensler, L. S.; Zhou, X.; Garchon, H. J.; Chiocchia, G.; Nossent, J.; Lie, B. A.; Forre, O.; Tuomilehto, J.; Laiho, K.; Bradbury, L. A.; Elewaut, D.; Burgos-Vargas, R.; Stebbings, S.; Appleton, L.; Farrah, C.; Lau, J.; Haroon, N.; Mulero, J.; Blanco, F. J.; Gonzalez-Gay, M. A.; Lopez-Larrea, C.; Bowness, P.; Gaffney, K.; Gaston, H.; Gladman, D. D.; Rahman, P.; Maksymowych, W. P.; Crusius, J. B.; van der Horst-Bruinsma, I. E.; Valle-Onate, R.; Romero-Sanchez, C.; Hansen, I. M.; Pimentel-Santos, F. M.; Inman, R. D.; Martin, J.; Breban, M.; Wordsworth, B. P.; Reveille, J. D.; Evans, D. M.; de Bakker, P. I.; Brown, M. A. Major Histocompatibility Complex Associations of Ankylosing Spondylitis Are Complex and Involve Further Epistasis with ERAP1. Nat Commun 2015, 6, 7146. https://doi.org/10.1038/ncomms8146.

(48) Alvarez-Navarro, C.; Lopez de Castro, J.A. ERAP1 Structure, Function and Pathogenetic Role in Ankylosing Spondylitis and Other MHC-Associated Diseases. Mol Immunol 2014, 57 (1), 12–21. https://doi.org/10.1016/j.molimm.2013.06.012.

(49) Stratikos, E.; Stamogiannos, A.; Zervoudi, E.; Fruci, D. A Role for Naturally Occurring Alleles of Endoplasmic Reticulum Aminopeptidases in Tumor Immunity and Cancer Predisposition. Front. Oncol. 2014. https://doi.org/10.3389/fonc.2014.00363.

(50) Serwold, T.; Gonzalez, F.; Kim, J.; Jacob, R.; Shastri, N. ERAAP Customizes Peptides for MHC Class I Molecules in the Endoplasmic Reticulum. Nature 2002, 419 (6906), 480–483. https://doi.org/10.1038/nature01074.

(51) Saric, T.; Chang, S. C.; Hattori, A.; York, I. A.; Markant, S.; Rock, K. L.; Tsujimoto, M.; Goldberg, A. L. An IFN-Gamma-Induced Aminopeptidase in the ER, ERAP1, Trims Precursors to MHC Class I-Presented Peptides. Nat Immunol 2002, 3 (12), 1169–1176.

(52) Hammer, G. E.; Gonzalez, F.; James, E.; Nolla, H.; Shastri, N. In the Absence of Aminopeptidase ERAAP, MHC Class I Molecules Present Many Unstable and Highly Immunogenic Peptides. Nat Immunol 2007, 8 (1), 101–108.

(53) Keller, M.; Ebstein, F.; Bürger, E.; Textoris-Taube, K.; Gorny, X.; Urban, S.; Zhao, F.; Dannenberg, T.; Sucker, A.; Keller, C.; Saveanu, L.; Krüger, E.; Rothkötter, H. J.; Dahlmann, B.; Henklein, P.; Voigt, A.; Kuckelkorn, U.; Paschen, A.; Kloetzel, P. M.; Seifert, U. The Proteasome Immunosubunits, PA28 and ER-Aminopeptidase 1 Protect Melanoma Cells from Efficient MART-126-35-Specific T-Cell Recognition. Eur. J. Immunol. 2015, 45 (12), 3257–3268. https://doi.org/10.1002/eji.201445243.

(54) James, E.; Bailey, I.; Sugiyarto, G.; Elliott, T. Induction of Protective Antitumor Immunity through Attenuation of ERAAP Function. J. Immunol. 2013, 190 (11), 5839– 5846. https://doi.org/10.4049/jimmunol.1300220.

(55) Barnea, E.; Melamed Kadosh, D.; Haimovich, Y.; Satumtira, N.; Dorris, M. L.; Nguyen, M. T.; Hammer, R. E.; Tran, T. M.; Colbert, R. A.; Taurog, J. D.; Admon, A. The Human Leukocyte Antigen (HLA)-B27 Peptidome in Vivo, in Spondyloarthritis-Susceptible HLA-B27 Transgenic Rats and the Effect of Erap1 Deletion. Mol. Cell. Proteomics 2017, 16 (4), 642–662. https://doi.org/10.1074/mcp.M116.066241.

(56) Koumantou, D.; Barnea, E.; Martin-Esteban, A.; Maben, Z.; Papakyriakou, A.; Mpakali, A.; Kokkala, P.; Pratsinis, H.; Georgiadis, D.; Stern, L. J. L. J.; Admon, A.; Stratikos, E. Editing the Immunopeptidome of Melanoma Cells Using a Potent Inhibitor of Endoplasmic Reticulum Aminopeptidase 1 (ERAP1). Cancer Immunol. Immunother. 2019, 68 (8). https://doi.org/10.1007/s00262-019-02358-0.

(57) Komov, L.; Kadosh, D. M.; Barnea, E.; Milner, E.; Hendler, A.; Admon, A. Cell Surface MHC Class I Expression Is Limited by the Availability of Peptide-Receptive “Empty” Molecules Rather than by the Supply of Peptide Ligands. Proteomics 2018, 18 (12), e1700248. https://doi.org/10.1002/pmic.201700248.

(58) Stratikos, E.; Stern, L. J. Antigenic Peptide Trimming by ER Aminopeptidases-Insights from Structural Studies. Mol. Immunol. 2013, 55 (3–4). https://doi.org/10.1016/j.molimm.2013.03.002.

(59) Ewer, K. J.; Barrett, J. R.; Belij-Rammerstorfer, S.; Sharpe, H.; Makinson, R.; Morter, R.; Flaxman, A.; Wright, D.; Bellamy, D.; Bittaye, M.; Dold, C.; Provine, N. M.; Aboagye, J.; Stafford, E.; et al. T Cell and Antibody Responses Induced by a Single Dose of ChAdOx1 NCoV-19 (AZD1222) Vaccine in a Phase 1/2 Clinical Trial. Nat. Med. 2021, 27 (2), 270–278. https://doi.org/10.1038/s41591-020-01194-5.

(60) Sahin, U.; Muik, A.; Derhovanessian, E.; Vogler, I.; Kranz, L. M.; Vormehr, M.; Baum, A.; Pascal, K.; Quandt, J.; Maurus, D.; Brachtendorf, S.; Lörks, V.; Sikorski, J.; Hilker, R.; Becker, D.; Eller, A. K.; Grützner, J.; Boesler, C.; Rosenbaum, C.; Kühnle, M. C.; Luxemburger, U.; Kemmer-Brück, A.; Langer, D.; Bexon, M.; Bolte, S.; Karikó, K.; Palanche, T.; Fischer, B.; Schultz, A.; Shi, P. Y.; Fontes-Garfias, C.; Perez, J. L.; Swanson, K. A.; Loschko, J.; Scully, I. L.; Cutler, M.; Kalina, W.; Kyratsous, C. A.; Cooper, D.; Dormitzer, P. R.; Jansen, K. U.; Türeci, Ö. COVID-19 Vaccine BNT162b1 Elicits Human Antibody and TH1 T Cell Responses. Nature 2020, 586 (7830), 594–599. https://doi.org/10.1038/s41586-020-2814-7.

(61) Reynolds, C. J.; Pade, C.; Gibbons, J. M.; Butler, D. K.; Otter, A. D.; Menacho, K.; Fontana, M.; Smit, A.; Sackville-West, J. E.; Cutino-Moguel, T.; Maini, M. K.; Chain, B.; Noursadeghi, M.; Brooks, T.; Semper, A.; Manisty, C.; Treibel, T. A.; Moon, J. C.; Valdes, A. M.; McKnight, Á.; Altmann, D. M.; Boyton, R. Prior SARS-CoV-2 Infection Rescues B and T Cell Responses to Variants after First Vaccine Dose. Science (80-.). 2021, eabh1282. https://doi.org/10.1126/science.abh1282.

(62) Perez-Riverol, Y.; Csordas, A.; Bai, J.; Bernal-Llinares, M.; Hewapathirana, S.; Kundu, D. J.; Inuganti, A.; Griss, J.; Mayer, G.; Eisenacher, M.; Perez, E.; Uszkoreit, J.; Pfeuffer, J.; Sachsenberg, T.; Yilmaz, S.; Tiwary, S.; Cox, J.; Audain, E.; Walzer, M.; Jarnuczak, A. F.; Ternent, T.; Brazma, A.; Vizcaino, J. A. The PRIDE Database and Related Tools and Resources in 2019: Improving Support for Quantification Data. Nucleic Acids Res. 2019, 47 (D1), D442–D450. https://doi.org/10.1093/nar/gky1106.

